# Bifurcating Neurons In The Anterior Thalamic Nuclei

**DOI:** 10.1101/2021.08.21.457238

**Authors:** Y Pei, S Tasananukorn, M Wolff, JC Dalrymple-Alford

## Abstract

The anterior thalamic nuclei (ATN) form a nodal point within a distributed memory network. The conventional view of ATN function describes segregated efferents to different terminal regions. By contrast, we found that bifurcating neurons are common within the anteromedial nucleus (AM) of the ATN. A substantial proportion of AM neurons (∼36% within a region of interest) showed collateral projections when one of two retrograde neurotracers, Cholera Toxin Subunit B (CTB) conjugated to either Alexa Fluor® 488 or 594, was placed in the medial prefrontal cortex (mPFC) or dorsal subiculum (dSub). A marked degree of collateralization (∼20% AM neurons) was also found when neurotracers were placed in mPFC and caudal retrosplenial cortex (cRSC); about 10% showed collaterals when the cRSC was paired with either dSub or vHF; the fewest (6%) was found for mPFC paired with the ventral hippocampal formation (vHF). A generally similar range of percentages of bifurcating neurons was found in the adjacent nucleus reuniens (Re). Evidence that AM neurons project simultaneously to many distantly-located structures provides a new perspective on ATN function. These neurons would facilitate direct coordination among key neural structures to support memory and may explain the strong association between the ATN and diencephalic amnesia.

## Introduction

Extensive clinical and experimental evidence supports the conclusion that the anterior thalamic nuclei (ATN) make a significant contribution to memory function (Aggleton, 2008; Aggleton et al., 2016; Barnett et al., 2021; Carlesimo et al., 2011; Dalrymple-Alford et al., 2015; Harding et al., 2000; Kim et al., 2009; Kopelman, 2015; Liu et al., 2021; Maillard et al., 2021; Sweeney-Reed et al., 2021; Wolff et al., 2015). This influence is consistent with the ATN’s strong reciprocal connections with three distinct memory structures, namely the medial prefrontal cortex (mPFC), retrosplenial cortex (RSC) and the hippocampal formation (HF) (Aggleton et al., 2016; Bubb et al., 2017; Mathiasen et al., 2017; Nelson, 2021; Shibata, 1993a, 1993b; van Groen et al., 1999; van Groen and Wyss, 1990; van Groen and Wyss, 1995). The effects of ATN lesions are sometimes more pronounced than lesions that disrupt their individual input or output structures (Dumont et al., 2015; Hamilton and Dalrymple-Alford, 2021; Mitchell et al., 2018; Perry et al., 2018; Warburton et al., 1999). Together, this evidence supports the perspective that the ATN form a critical subcortical hub that actively integrates information processing among telencephalic memory structures (Wolff and Vann, 2019).

If the ATN region plays an active role that is vital for normal memory processing, then it is critical to understand its efferent projections. Currently, ATN neural projections to different memory-related regions are viewed as contributing predominantly to separate neuronal populations that give rise to hierarchically organised subsystems (Aggleton et al., 2010; Jankowski et al., 2013; Mathiasen et al., 2020; Nelson, 2021; Wright et al., 2013). This perspective emphasises functionally segregated ATN circuits associated with the anteromedial (AM), anteroventral (AV) and anterodorsal (AD) nuclei, with distinct albeit complementary contributions to spatial memory and mental navigation in particular (Aggleton and Nelson, 2015; Aggleton et al., 2010). It is clear that there are very few bifurcating afferents to the ATN from cortical structures, and convergence of parallel information streams within the ATN, but the thalamo-cortical projections that emanate from different ATN nuclei show regional overlap in many of their terminal fields (Aggleton et al., 2010; Mathiasen et al., 2020; Nelson, 2021; Wright et al., 2013). The idea that the ATN actively integrate different information streams among cortical structures (Wolff and Vann, 2019), however, would be significantly strengthened by evidence that a subpopulation of ATN neurons send collateral projections to distinct terminal regions, so as to influence the mPFC, RSC and / or HF simultaneously.

Current descriptions of AM connections suggest that this nucleus in particular may be a region with bifurcating neurons. The AM is the primary source of ATN efferents to the mPFC. We also know that the AM provides input to the RSC and HF, being the prominent source of ATN efferents to the subiculum (Bubb et al., 2017; Jankowski et al., 2013; Shibata, 1993a, 1993b; van Groen and Wyss, 1995; van Groen et al., 1999). We therefore asked whether a subpopulation of AM neurons have a broad influence on the memory system by sending axon collaterals to distantly-located pairs of memory-related structures. The retrograde neurotracer Cholera Toxin Subunit B (CTB), conjugated to either Alexa Fluor® 488 or 594, was used to determine if bifurcating neurons exist for five pairs of terminal regions. Three neurotracer pairings focused on the mPFC when paired with either the dorsal subiculum (dSub), ventral HF (vHF), or caudal RSC (cRSC). Two neurotracer pairings examined if AM neurons send collaterals to the cRSC when paired with either the dSub or vHF. We used CTB conjugates as these are sensitive and reliable retrograde neurotracers (Conte et al., 2009a, 2009b) that have successfully identified double-labelled neurons in the adjacent thalamic reuniens (Re) immediately ventral to the AM (Varela et al., 2014). We found a surprisingly marked degree of AM neuron collateralizations, especially when one retrograde neurotracer was infused in the mPFC and the alternate neurotracer was placed in the dSub or cRSC.

## Results

### Hemispheric expression of neurotracer labelling

Uptake of the neurotracers in ATN neurons, in all cases, was almost exclusively ipsilateral to the hemisphere in which the infusions were made. In all cases (i.e., after tracer infusions in all the four terminal field structures), labelling was also found in the Re, which rarely crossed the midline. Unlike the ATN and the Re, there was some contralateral uptake in the mediodorsal thalamic nucleus associated with the mPFC infusion and some uptake of neurotracer across the midline region of the paraventricular thalamic nucleus associated with the vHF infusion. In both of these cases, however, the contralateral expression was much weaker than that shown on the ipsilateral side.

### Placement of neurotracers and uptake in bifurcating neurons

Fourteen cases had correctly-placed infusions of the retrograde tracers in both target structures (Fig. 1). Fig. 1 also shows the location of the infusions made in case Y6 (excluded due to misplaced vHF infusion). Evidence from counterbalanced infusions of the specific tracers at the two sites for any given paring of neurotracers showed the same pattern of findings. Double-labelled neurons in the AM were found for each of the five neurotracer location pairings (Table 1). Cell counting in the AM region (Fig. 2) is described below.

**Table 1.**
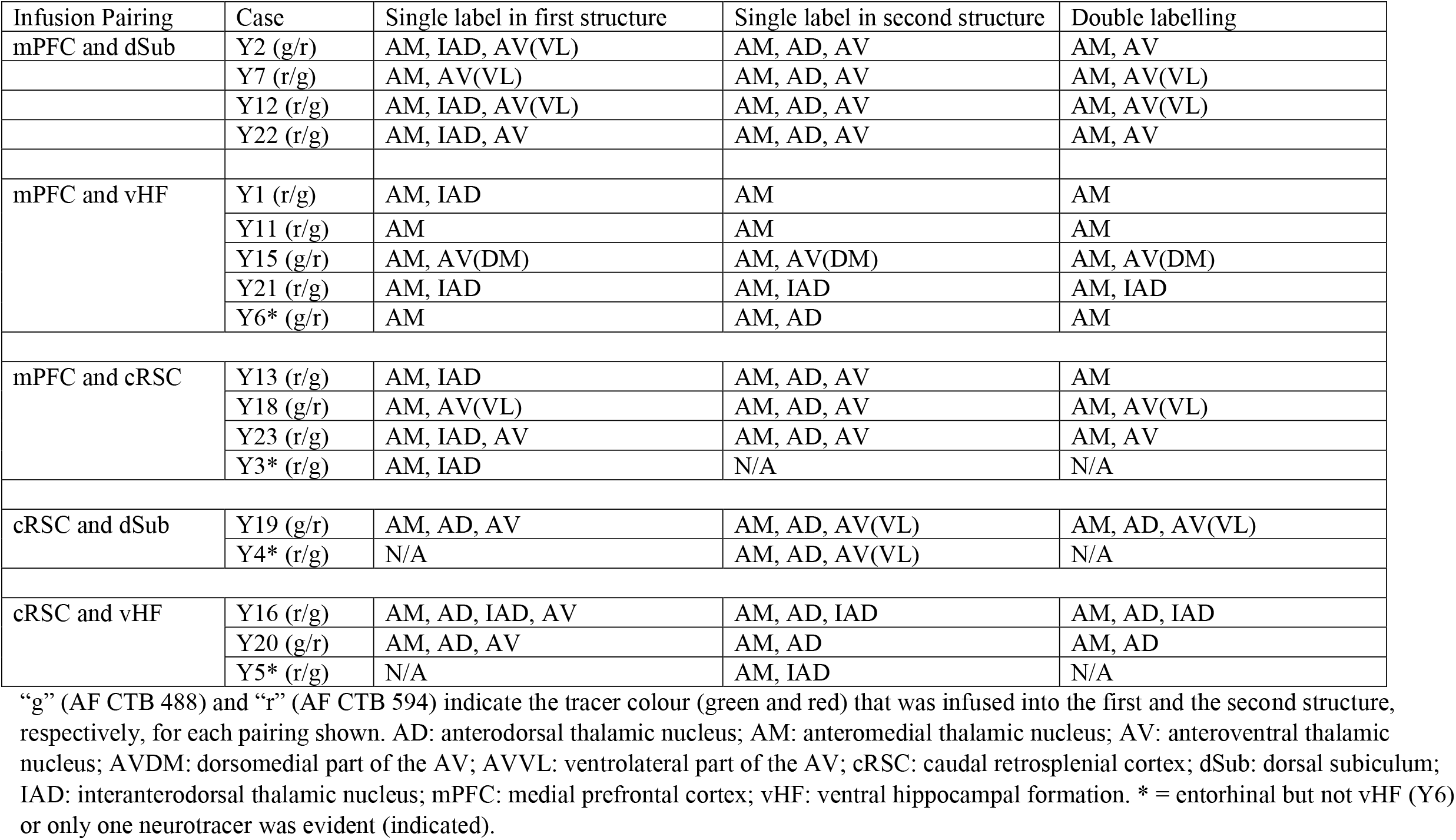
Areas in the ATN that show single and double-labelled cells for each rat after infusion of the retrograde tracers (AF CTB 488 and AF CTB 594) into a pair of structures for five different pairings.

**Fig. 1.**
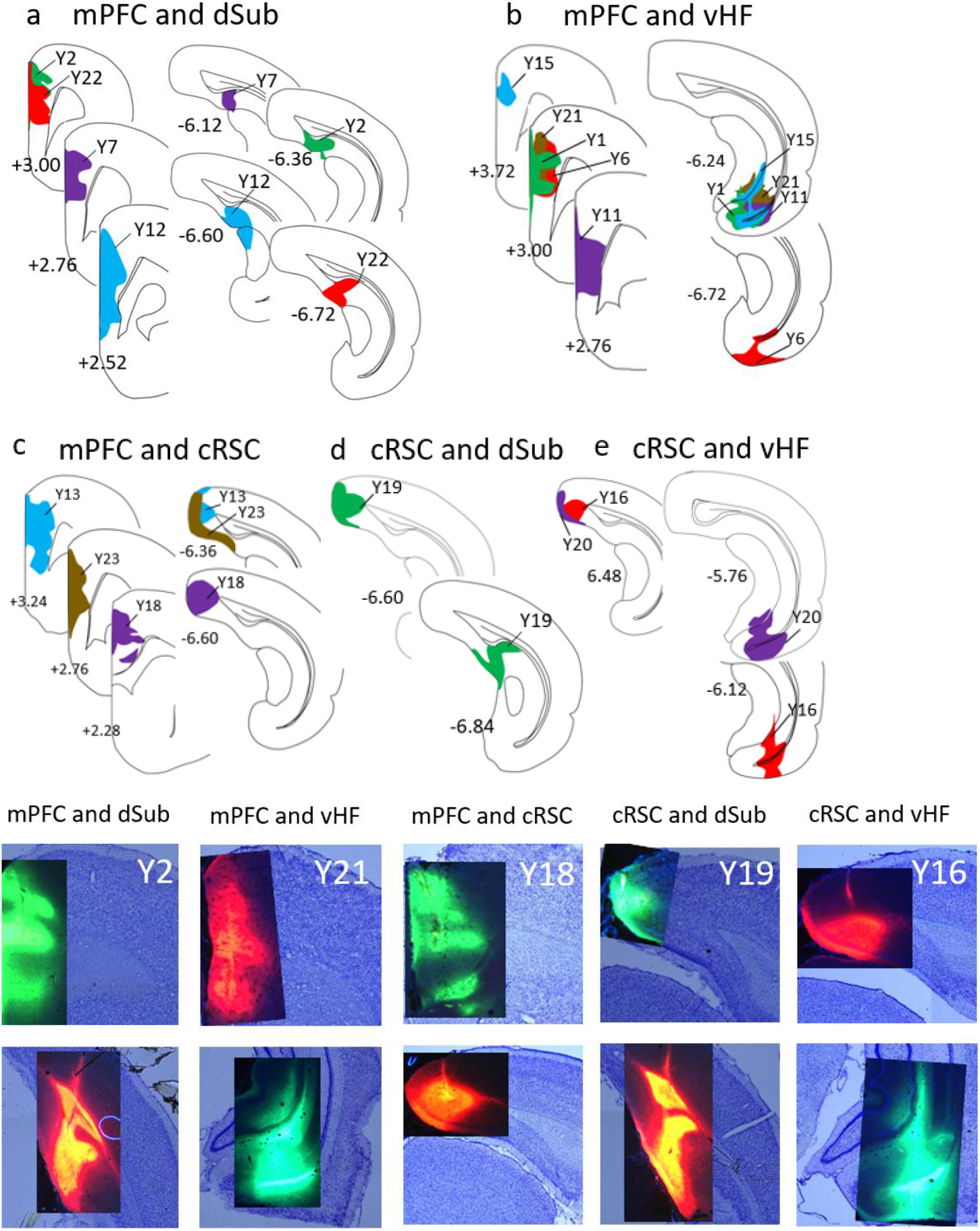
Upper panels show schematic representations of paired infusions of neurotracers AF CTB 488 and AF CTB 594 for 15 individual rats in the five paired structures used. Within each terminal-field pairing, the same rat is shown using the same colour and different rats are shown with different colours. The lower pair of rows show photomicrographs of the infusions, overlaid on the same section that was counterstained with cresyl violet, for one representative rat per neurotracer pairing. The upper row of this pair shows the infusion in the first of the paired structures and the lower row shows the corresponding infusion in the second terminal field. Note that none of the infusions spread to the contralateral hemisphere. AF CTB: Cholera Toxin Subunit B (Recombinant) Alexa Fluor™; cRSC: caudal retrosplenial cortex; dSub: dorsal subiculum; mPFC: medial prefrontal cortex; vHF: ventral hippocampal formation.

**Fig. 2.**
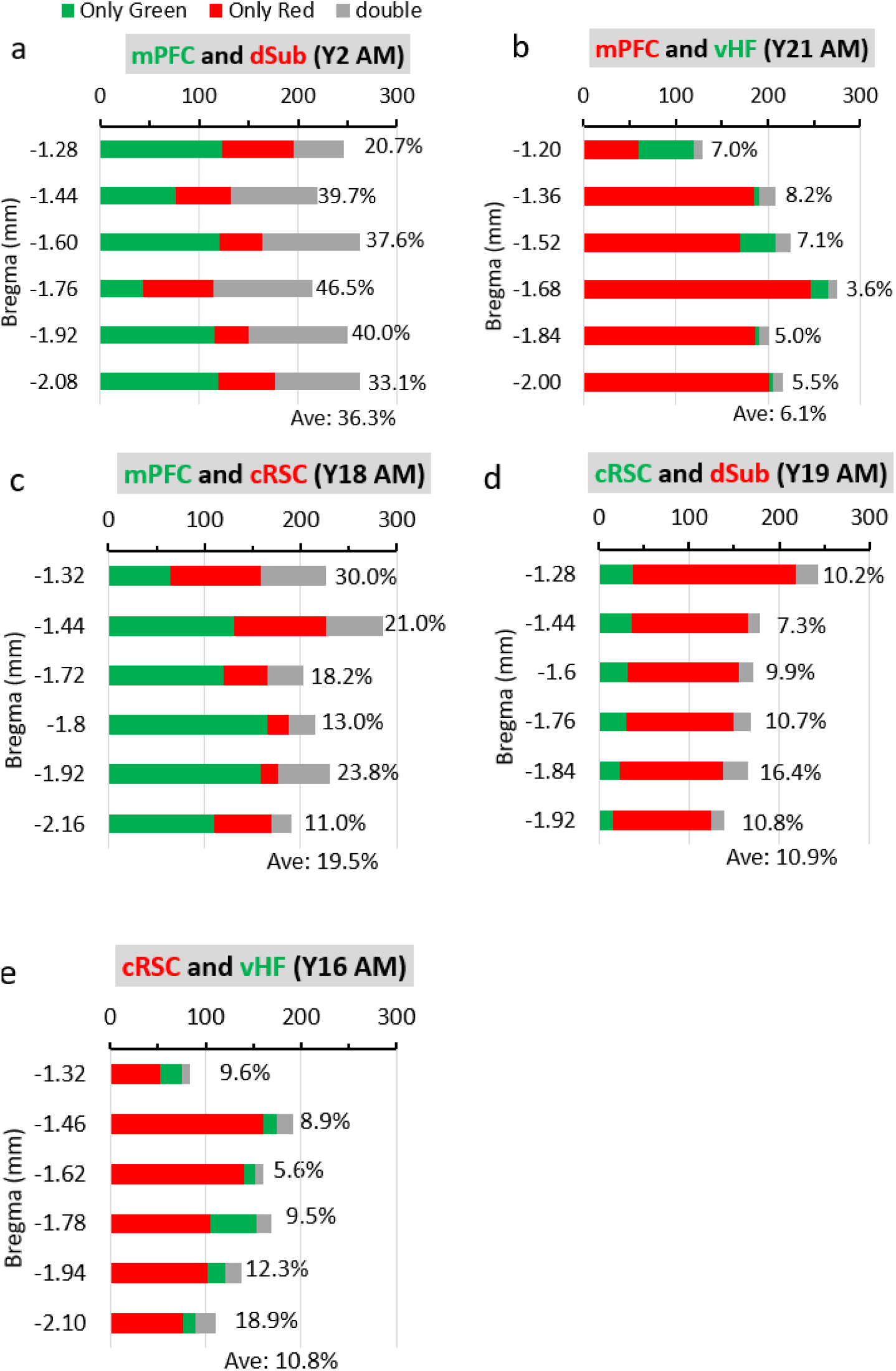
Number and relative percentage of labelled cells in the AM at six rostral-caudal levels in a representative rat for each of the five neurotracer pairings. The percentage of bifurcating neurons (grey bars) is expressed relative to the total number of neurons identified by both tracers (including bifurcating ones). The number of cells showing just one label excluded all bifurcating cells. Infusions were made with AF CTB 488 (green) in one structure and AF CTB 594 (red) in the other structure. The colours used to label structures above each panel correspond to the tracer used for that structure. Ave: average percent of double labelled cells across the six AM levels. AF CTB: Cholera Toxin Subunit B (Recombinant) Alexa Fluor™; AM: anteromedial thalamic nucleus; cRSC: caudal retrosplenial cortex; dSub: dorsal subiculum; mPFC: medial prefrontal cortex; vHF: ventral hippocampal formation.

#### mPFC and dSub

These rats were Y2, Y7, Y12, and Y22 (Fig. 1). The mPFC infusion was centred in the dorsal and mid-ventral aspects of the mPFC, including A32D and A32V in both Y2 and Y22, and A24b and A24a in both Y7 and Y12. Note that A32D and A24b are generally equivalent to the cingulate cortex area 1 (Cg1) and A32V and A24a include much of what is classically regarded as prelimbic cortex (PrL) (see Vogt and Paxinos, 2014). There was additional posterior and ventral spread in the mPFC in Y12, at A33 and A25, generally equivalent to PrL and infralimbic cortex (IL), respectively. The dSub infusion included the subiculum in all cases and the postsubiculum in all cases except Y7. Neurotracer was also present in the presubiculum in Y22.

As shown in Table 1, the pattern of labelled cells in the thalamus was consistent across all four rats. The mPFC infusion produced dual-labelled neurons predominantly in the AM, which was most prominent in the lateral and dorsal regions (Fig. 3). There was some dual-labelling in the AV, usually in the ventrolateral part of the AV (AVVL). Single labelling after the dSub infusion was associated with neurons located in all three nuclei (AM, AV and AD), but most strongly in AV and AD regions.

**Fig. 3.**
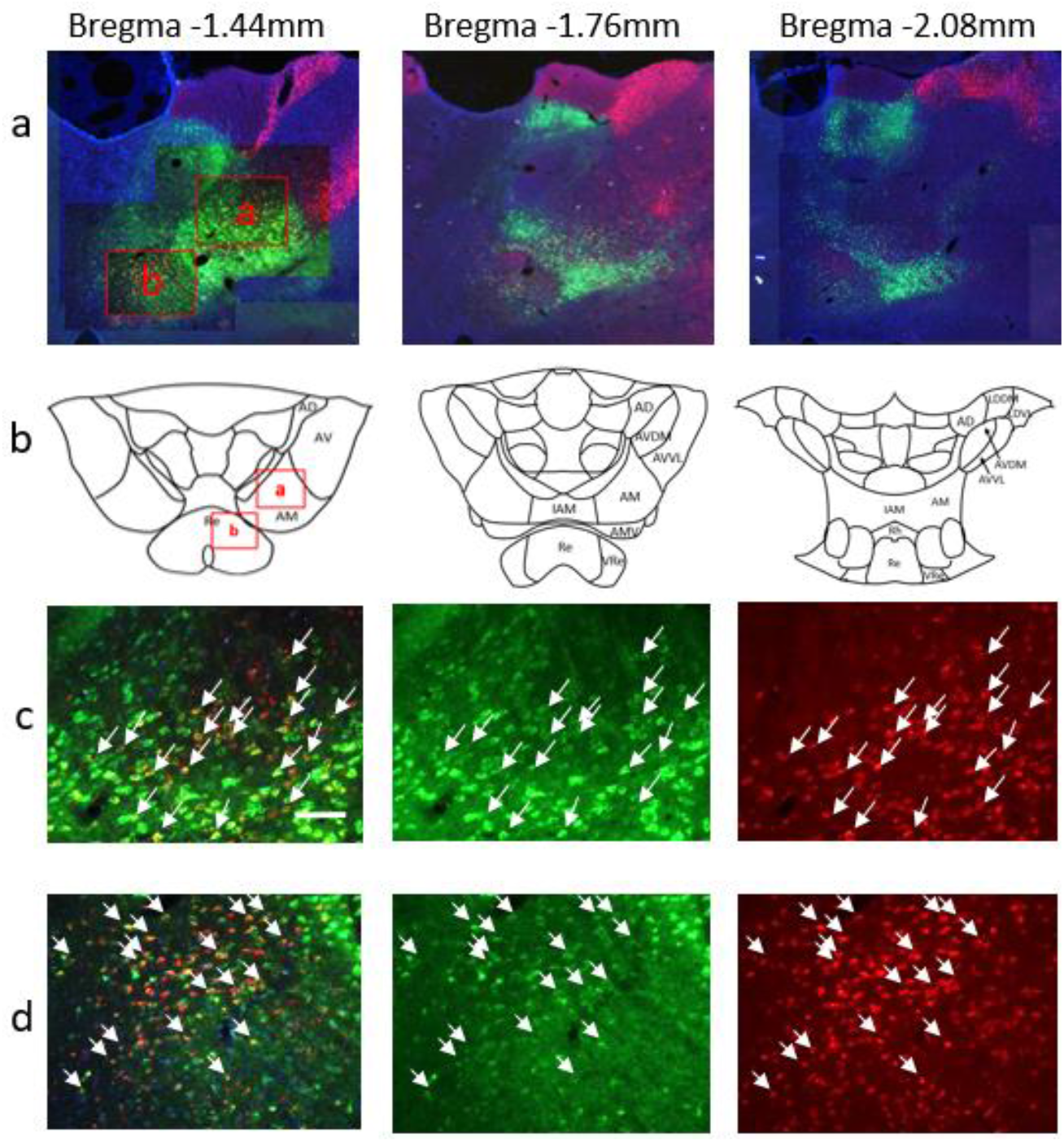
Single and double labelled cells in the AM and Re at three rostral-caudal levels in Y2 after infusion of AF CTB 488 (green) into mPFC and AF CTB 594 (red) into dSub. The position of the red-box insets in the first panel of row (a) are depicted on the atlas outline in row (b) at Bregma - 1.44 mm. (c) Higher magnification view of inset box a within the top left panel, showing the AM, with examples of double-labelled cells indicated by white arrows in the left panel and corresponding green and red labelled cells in the middle and right panels. Row (d) shows the corresponding fluorescence but now in the inset box b from row (a), which is the Re region. Scale bar 100um in c. AD: anterodorsal thalamic nucleus; AF CTB: Cholera Toxin Subunit B (Recombinant) Alexa Fluor™; AM: anteromedial thalamic nucleus; AMV: anteromedial thalamic nucleus, ventral part; AV: anteroventral thalamic nucleus; AVDM: dorsomedial part of the AV; AVVL: ventrolateral part of the AV; IAM: interanteromedial thalamic nucleus; LDDM: laterodorsal thalamic nucleus, dorsomedial part; LDVL: laterodorsal thalamic nucleus, ventrolateral part; Re: reuniens thalamic nucleus; Rh: rhomboid thalamic nucleus; vRe: ventral reuniens thalamic nucleus.

#### mPFC and vHF

These rats were Y1, Y11, Y15, and Y21 (Fig. 1). The mPFC infusion was centred at A32D and A32V in four cases. The infusion in Y11 was more posterior and ventral, spreading to A24a and A25. The vHF infusion was centred in the ventral subiculum in four cases and included entorhinal cortex in all cases. There was some spread into the molecular layer of the dentate gyrus (MoDG) in cases Y15 and Y21.

Bifurcating cells from infusions in mPFC and in vHF were again almost exclusively found in the AM in all 4 rats. There were some bifurcating neurons in the dorsomedial part of the AV (AVDM) in rat Y15 and some in the interanterodorsal thalamic nucleus (IAD) in rat Y21 (Table 1). Single projecting cells were almost exclusively found in the AM in all four rats. Case Y6, with an infusion in only the entorhinal cortex and not the vHF, also showed some double-labelled cells in the AM.

#### mPFC and cRSC

These cases were Y13, Y18 and Y23 (Fig. 1). The mPFC infusion was evident in both dorsal and ventral mPFC in Y13, covering A32D, A32V, and A25. The mPFC infusion was more posterior in the other two cases and was centred in A24b and A24a. In all cases, the cRSC infusions covered A30, A29c, and A29b, with additional spread in A29a in Y23. Note that A30 is generally known as the retrosplenial dysgranular cortex (RSD), and A29c, A29b, and A29a are known as the retrosplenial granular cortex c region (RSGc), retrosplenial granular cortex b region (RSGb), and retrosplenial granular cortex a region (RSGa), respectively.

Evidence of bifurcating neurons was found in the AM in all three cases. There were bifurcating cells also for the AV in cases Y18 and Y23. In all three cases, single-projecting cells from the infusions in the mPFC were again located in the AM with less evidence of single-labelled cells in the AV. A few cells labelled by mPFC infusions were scattered in the IAD in Y13 and Y23. Single-labelled cells from the cRSC infusions were clearly evident in the AD and AV regions in all three rats, but there were also single-labelled cells in the AM. In the fourth case, Y3, the cRSC infusion failed and single-labelled cells from the mPFC infusion were found in the AM and IAD.

#### cRSC and dSub

There was one case (Case Y19), which had a cRSC infusion that included A30, A29c, and A29b and a dSub infusion that included the subiculum, the postsubiculum, and presubiculum (Fig. 1). Both double-labelled and single-labelled cells were evident across all three ATN nuclei. Double labelled cells were more frequently found in the AM and in the region where the AVDM intersects with the AD and in the AVVL. The single-labelled cells were more frequent in the AVDM after cRSC infusion, and in the AD, AM and AVVL following dSub infusion. The second case, Y4, only had a successful infusion in the dSub (the cRSC infusion failed in this rat). For Y4, single-labelled cells from the dSub infusion were found in all three ATN nuclei, similar to the single-labelled cells found for Y19.

#### cRSC and vHF

These cases were Y16 and Y20 (Fig. 1). The cRSC infusion was located in A29c and A29b for both rats, with some tracer in A29a for Y20. Both vHF infusions included the ventral subiculum with additional spread into the entorhinal cortex in Y16 and in the MoDG in Y20.

Bifurcating cells were found in the AM and to a lesser extent in the AD in both cases; case Y16 also showed double-labelled cells in the IAD. The cRSC infusion labelled cells in the AM, AV and AD for both cases. The vHF infusion produced labelled cells in the AM and AD for both cases. Case Y16 showed single-labelled cells from both infusions in the IAD. The third case, Y5, had a failed infusion in the cRSC and showed single labelled cells from the vHF infusion in the AM and IAD.

#### Cell counts in the AM

Regions of interest within the AM were examined, based on initial observation of tissue samples. Estimates of the number of single-labelled and double-labelled AM neurons for a representative rat for each of the five neurotracer pairings are shown in Fig. 2. A photomicrograph example of the bifurcating AM neurons after neurotracer infusions in the mPFC and dSub (Y2) is shown in Fig. 3.

For the mPFC and dSub pairing (Y2), the proportion of bifurcating neurons in the AM was substantial (36% overall) and consistently high throughout this nucleus (Fig. 2a). This proportion rose from 20% for the most anterior section (at Bregma -1.28 mm) to about 40% for the remainder of the AM except for the most posterior section where it fell to 33% (at Bregma -2.08 mm). The fraction of double labelled cells was slightly higher in the dSub-projecting population than in the mPFC-projecting population (average 61% and 48%, respectively).

There was also a marked proportion of bifurcating AM neurons (20% overall) for the mPFC and cRSC pairing (Y18). This was 30% for the most anterior AM region with generally lower proportions in photomicrographs from more caudal sections (Fig. 2c). A higher fraction of double-labelled cells was found in the cRSC-projecting cells (47%) compared to that in the mPFC-projecting cells (27%).

The proportion of AM bifurcating cells was much lower for the other three pairings, averaging 6% for mPFC and vHF, 11% for cRSC and dSub, and 11% for cRSC and vHF (Fig. 2). These proportions were generally consistent across the anterior to posterior extent of the AM. In the mPFC and vHF pairing, the double-labelled cells comprised almost half of the total subpopulation of vHF-projecting cells (50%) but far lower for that of the mPFC-projecting cells (7%), which was possibly related to the substantially lower total number of cells that projected to the vHF than projected to the mPFC. For the cRSC and dSub pairing, the fraction of double labelled cells was considerably higher in the cRSC-projecting cells (40%) than in the dSub-projecting cells (13%), perhaps now reflecting a much higher total number of dSub-projecting cells than cRSC-projecting cells. With respect to the cRSC and vHF pairing, the fraction of double-labelled cells was considerably higher in the vHF-projecting cells (43%) than in the cRSC-projecting cells (13%), which may be associated with the disproportionally higher total number of cRSC-projecting cells than vHF-projecting cells that was captured by the infusions made.

While the primary focus of the current study was the AM, we also examined some photomicrographs in regions of interest in the AV and AD subregions. Across the rostral extent of these regions, about 8% to 20% of double-labelled AV neurons were found for the mPFC and dSub pairing, 24-37% for mPFC paired with cRSC, 23% for the mPFC and vHF pairing, and 9-22% for the cRSC and dSub combination. No double-labelled cells in the AV were found when the cRSC was paired with the vHF. Photomicrographs for the AD suggested 8-31% double-labelled neurons for the cRSC paired with dSub, and 11% for the cRSC and vHF pairing, but none for any pairing that included the mPFC.

### Bifurcating neurons in the Re region ventral to the AM

In the same sections that were assessed for ATN cell labelling, single-labelled neurons were found in regions of interest in the Re after neurotracer infusions placed in all four terminal sites. Double-labelled Re neurons were also evident in all five neurotracer pairings (Fig. 4). As with the ATN, the largest proportion of double-labelled cells was found with the mPFC and dSub pairing in case Y2, which averaged 31% and was generally consistent throughout the Re to the extent that we examined the Re region ventral to the ATN. There was also a relatively high proportion of bifurcating cells in the Re associated with the mPFC and cRSC pairing (26%) in rat Y18. We were unable to collect all the intended anterior to posterior photomicrographs for the Re in the other three infusion pairings (mPFC and vHF; cRSC and dSub; cRSC and vHF), but the available evidence in Fig. 4 suggests perhaps more bifurcating Re neurons for the pairing mPFC with vHF (11%) than was found for the AM for this pairing (6%), but a similar proportion of bifurcating neurons for cRSC with dSub (12%) and for cRSC with vHF (14%) in Re as was found for AM (11% in both cases).

**Fig. 4.**
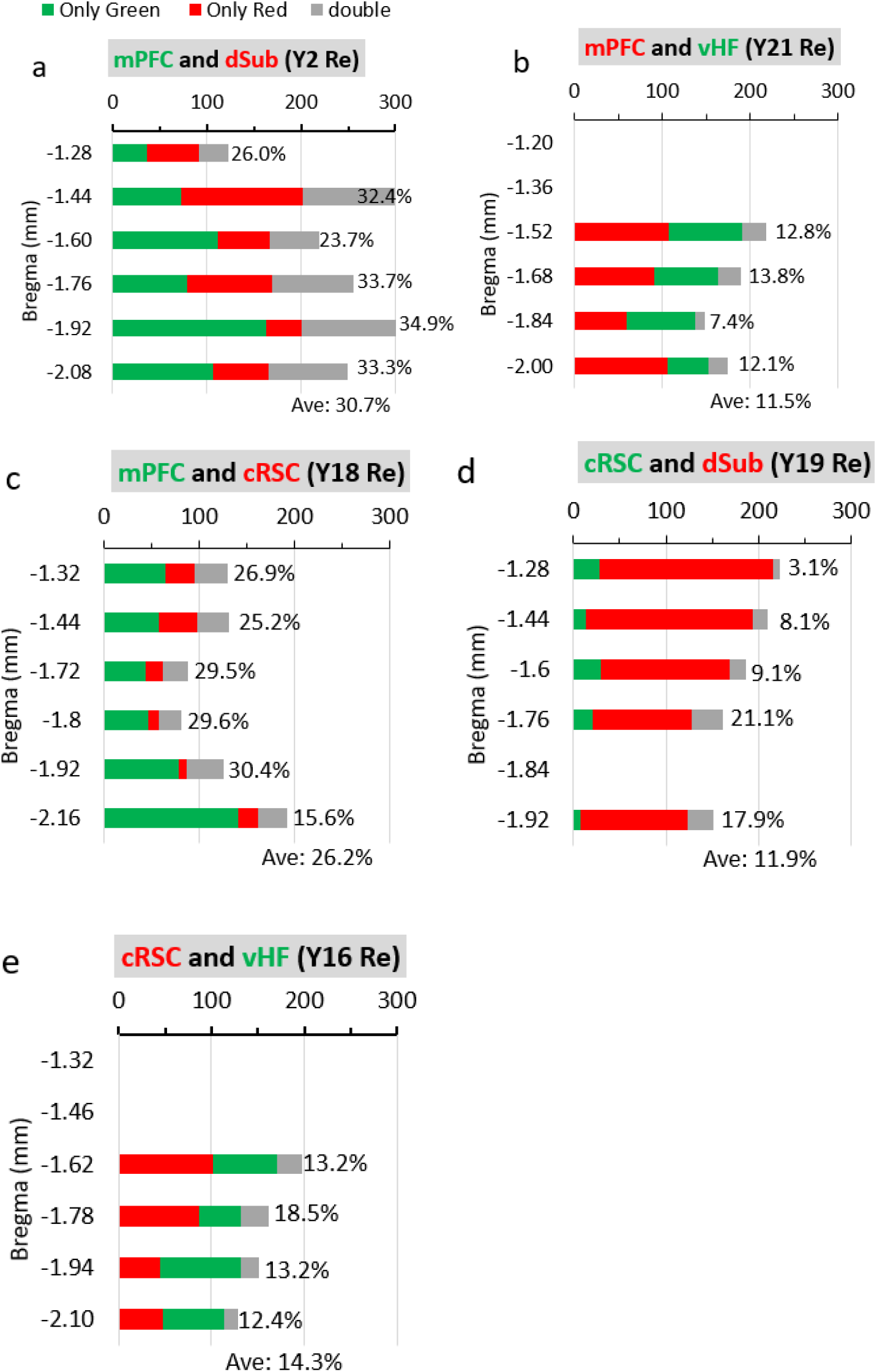
Number and relative percentage of labelled cells in the Re across rostral-caudal levels in a representative rat for each of the five neurotracer pairings. The percentage of bifurcating neurons is expressed relative to the total number of neurons identified by both tracers (including bifurcating ones). The number of cells showing just one label excluded all bifurcating cells. Infusions were made with CTB 488 (green) in one structure and CTB 594 (red) in the other structure. The colours used to label structures above each panel correspond to the tracer used for that structure. Photomicrographs were not taken at levels where no bar is shown. Ave: average percent of double labelled cells across all the available levels of Re for that rat; AF CTB: Cholera Toxin Subunit B (Recombinant) Alexa Fluor™; cRSC: caudal retrosplenial cortex; dSub: dorsal subiculum; mPFC: medial prefrontal cortex; Re: reuniens thalamic nucleus; vHF: ventral hippocampal formation.

In the Re, the mPFC and dSub pairing, the mPFC and vHF pairing, and the cRSC and vHF pairing all produced similar fraction of double-labelled cells relative to the total population that projected to each of the former and latter sites within each pair of infusions (45% and 52% for mPFC and dSub, respectively; 19% and 24% for mPFC and vHF, respectively; 25% and 27% for cRSC and vHF, respectively).

For the mPFC and cRSC pairing, however, the proportion of double labelled cells in the Re was considerably higher in the cRSC-projecting population (61%), being almost double that of the mPFC-projecting population (32%). Similarly, for the cRSC and dSub pairing, the double-labelled cells comprised a significantly higher proportion of the cRSC-projecting population (50%) compared to the dSub-projecting population of cells (13.2%) in the Re, possibly due to the much higher total number of cells labelled by the dSub infusion than by the cRSC infusion.

## Discussion

We used a double retrograde CTB fluorescent tracing technique to examine whether there are bifurcating neurons in the ATN that project to separate, distantly-located cortical regions. CTB neurotracers overcome the potential imbalance of labelling when different retrograde tracers are used (e.g., Horikawa et al., 1988; Hoover and Vertes, 2012). Our approach revealed a high number of neurons in the AM nucleus that send collaterals to multiple memory-related brain regions, including mPFC, RSC and HF. Bifurcating neurons were found for all five of the neurotracer pairings examined, specifically, mPFC paired with either dSub, vHF, or cRSC, and cRSC paired with either dSub or vHF. Although not examined in detail, some bifurcating neurons were also evident in the AV and AD nuclei of the ATN, so more work is required to establish collateralizations from these subregions. Previously, the prospect of bifurcating ATN neurons was examined only from the perspective of adjacent regions in the cingulate gyrus, which produced mixed results (Sripanidkulchai and Wyss, 1986; Horikawa et al., 1988). Consistent with our findings, however, Horikawa and colleagues reported that about 9% to 13% of the projection neurons from the AM nucleus send bifurcating axons when more distant subregion pairs of the cingulate cortex were examined, that is, the rostral ACC and RSC (Horikawa et al., 1988). Like previous descriptions of single ATN efferent projections (Mathiasen et al., 2017), the bifurcating ATN efferents that we observed were also exclusively unilateral. Previous single-label neurotracer studies have shown that the ATN have multiple efferents that can exert an influence across an extended memory system (Bubb et al., 2017; Shibata, 1993b, 1993a; van Groen et al., 1999; van Groen and Wyss, 1995). Our new evidence that AM neurons show a marked degree of collateralization to many distantly-located cortical structures supports a conceptual advance on current ideas of ATN function.

The standard view of the ATN’s role in a hippocampal-diencephalic-cortical memory network emphasises distinct neural circuits that broadly distinguish the AM, AV and AD nuclei and their connections (Aggleton et al., 2010; Jankowski et al., 2013; Nelson, 2021; Wright et al., 2013). This perspective is predicated on a system of non-bifurcating ATN efferents that project, separately, to other neocortical and limbic cortex structures. While each component of the ATN exhibits different output mappings, our evidence suggests that these distinctions may be less prominent than previously envisaged. Indeed, the relative abundance of bifurcating neurons in the AM, with collaterals to many of the different terminal regions in the extended memory system, has not been anticipated by prior models. This suggests that the AM in particular plays a significant role in the integration of memory-related information processing across the extended memory system. The functional relevance of these bifurcating neurons may be similar to that posited for bifurcating neurons in the Re (Hoover and Vertes, 2012; Varela et al., 2014). That is, AM bifurcating neurons, and in particular the high number of cells sending collaterals to both mPFC and dSub, may facilitate the coordination and synchronization of interactions across the prefrontal cortex and hippocampal regions, two key memory regions in the brain (Eichenbaum, 2017).

Our findings make a significant contribution to the literature supporting a critical role for the ATN in memory. The wide range of AM neuron collateralizations would support a significant degree of neural integration across the memory system. These collateralizations could play a unique role in maintaining and updating mental representations of spatial and nonspatial information in the manner proposed by Wolff and Vann (2019). In support of this, electrophysiological recordings made in drug-resistant epilepsy patients suggest that the ATN integrate information from diverse cortical sources during encoding to guide successful recall (Sweeney-Reed et al., 2021). Other recent studies have shown that working memory in both humans and rats can be improved by stimulation of the ATN (Barnett et al., 2021; Liu et al., 2021). It would be especially informative to learn if the bifurcating ATN neurons have a pre-eminent role in enhancing memory. Experimental manipulations will be needed that selectively target these various bifurcating neurons to determine their role in the functional organisation of the ATN. The discovery of bifurcating neurons in the ATN may help answer why injury or dysfunction of this region is strongly associated with diencephalic amnesia and other disorders associated with poor memory (Aggleton et al., 2016; Carlesimo et al., 2011; Harding et al., 2000; Perry et al., 2019).

We found that AM neurons send axons to all four terminal regions examined (i.e. mPFC; dSub; vHF; and cRSC), but the proportion of double-labelled neurons was not the same across the five neurotracer pairings. The highest proportion (36%; range 21-47% across rostro-caudal regions of interest) was found after infusions in the mPFC and dSub terminal regions. It should be noted that we did not look systematically across the whole of the AM nucleus, so the true proportion may vary once additional follow-up studies are completed. Nonetheless, comparisons across the different pairings suggested far lower proportions of bifurcating AM neurons (6-11%) after pairing the mPFC with vHF, and cRSC with either dSub or vHF. An intermediate but marked proportion of bifurcating AM neurons was found after neurotracer infusions were placed in the mPFC and cRSC (20%). The proportion of double-labelled cells relative to the corresponding single-labelled cells also varied. This proportion was higher for the dSub-projecting neurons when compared to the proportion projecting to the mPFC, but it was lower for the dSub-projecting neurons when compared to the proportion projecting to the cRSC. With respect to the vHF-projecting neurons, the proportion of double-labelled cells relative to the single labelled projections was higher than was the case for the corresponding mPFC and cRSC projections. Our findings may reflect the relatively large neurotracer volumes used, which was to avoid the potential issue of under-estimating the presence of bifurcating cells with small volumes of neurotracer. Our infusions remained within the relevant target structures, but we did not make a systematic comparison across the whole of each terminal region. Nonetheless, our findings suggest a different magnitude of influence by bifurcating neurons across the different terminal regions examined. We might also expect the proportions found to vary with the topography of each terminal region. Estimates derived from infusions across a wider area of the subiculum would reinforce the full extent of collaterals from ATN neurons, and specifically AM neurons, in this important hippocampal structure. It would also be interesting to examine additional regions, especially more caudal aspects of the ACC and the more rostral aspects of the RSC. Another issue is that we have only looked at collateralizations for pairs of ATN efferent targets. It is possible that some neurons have a more widespread influence by projecting to more than two target structures, which has been found to be the case for mediodorsal thalamus neurons (Kuramoto et al., 2017).

Although our focus was the AM subregion of the ATN, we also provide evidence on bifurcating neurons in the area of the rostral Re that is ventral to the AM. Like previous studies on the Re, pairing two different neurotracers across the mPFC and vHF revealed a modest proportion of bifurcating Re cells (Hoover and Vertes, 2012; Varela et al., 2014). This consistency with prior work on the Re is important as it increases confidence in our overall methods and findings. For our study, the proportion of dual-labelled Re neurons for the mPFC and vHF pairing was 11.5%; previous studies reported about 3-9% of bifurcating Re cells, but the average was higher (8%) when, like us, CTB tracers were used (Varela et al., 2014). One explanation of the small differences compared to our study is that the two previous Re studies used a smaller volume of neurotracer infusions in the mPFC and vHF. We used relatively large volumes of neurotracers, in our case in two sites per target, for the reasons given above. Another important difference was that prior work on the Re targeted the ventral CA1 region, which produced a larger number of bifurcating Re neurons compared to dorsal CA1 infusions, but they also tended to find more labelling when their ventral infusions included the subiculum region. By contrast, our infusions explicitly targeted the ventral subiculum and avoided CA1. Our findings on bifurcating neurons in the Re extend these earlier studies, as we provide new evidence that Re neurons, like AM neurons, send collaterals to many other brain regions. Specifically, we found a relatively high number of bifurcating Re neurons when the mPFC was paired with either dSub (30%) or cRSC (26%), and a moderate number when the cRSC was paired with either dSub (12%) or vHF (14%). One particularly interesting observation was that the proportion of bifurcating Re neurons was two or three-fold for the mPFC pairing with dSub or cRSC than was the case for mPFC paired with vHF, because this latter pairing has been the focus of previous studies which will therefore have underestimated the diverse manner in which the Re region can simultaneously impact memory-related structures. While these proportions must be treated with caution, and in the context of our methodology, they demonstrate a much wider influence of bifurcating Re neurons on the extended memory system than previously described. Other studies have found that the Re sends collaterals to both mPFC and other non-cortical regions such as the nucleus accumbens (see Cassel et al., 2013). Like the Re, it is possible that a wider range of target structures that receive collaterals needs also to be examined with respect to the AM region.

While both ATN and Re can be regarded as providing a neural substrate that mediates mPFC and hippocampal interactions, the general consensus is that these two thalamic regions have complementary roles in memory and other cognitive processes (Mathiasen, et al., 2020; Wolff and Vann, 2019). Neuroanatomy has been used to support this difference. The Re has dense connections with hippocampal CA1, especially ventral CA1 (Hoover and Vertes, 2012; Tao et al., 2021) whereas the ATN has dense connections with the subicular cortex (Bubb et al., 2017; Mathiasen et al., 2020; Nelson, 2021). Nonetheless, the Re sends efferents to the subiculum. Also, there is growing evidence that the ATN also has some projections to CA1 (de Lima et al., 2017; Prasad and Chudasama, 2013; Tao et al., 2021). In terms of memory function, not all lesion or inactivation studies are consistent, but the Re is most often associated with long-term memory consolidation rather than retrieval for spatial memory and contextual memory (de Vasconcelos and Cassel, 2015; Ferarris et al., 2021; Quet et al., 2020). ATN lesions more consistently disrupt performance on spatial working memory, and acquisition of reference spatial memory and contextual fear conditioning (Aggleton and Nelson, 2015; Marchand et al., 2014; Perry et al., 2018; Vetere et al., 2021). Nonetheless, it will be important to compare manipulations of the ATN and Re within the same task conditions and examine possible functional heterogeneity within each thalamic region. This may be especially informative when focusing on the AM as the comparison for Re because both regions include a significant number of neurons that send collaterals to broadly similar terminal regions.

Our study has limitations and needs to be assessed in the context of the methodology used. We detailed evidence on the existence of bifurcating neurons in the AM. More systematic work that is focused on the neurons with collateralizations emanating from the AD and AV regions is also warranted. Our method of analysis of the bifurcating neurons in the AM and Re was similar to previous work that focused on the Re (Hoover and Vertes, 2012) and relied on semi-quantitative analysis of dual excitation of neurons in the same focal plane. In our case, we took photomicrographs to look at regions of interest across the rostro-caudal extent but we did not cover the entire 2-dimensional boundary of the AM or Re, so our cell count and percent values must be treated with caution. Nonetheless, we were careful to count cells only when fluorescence surrounded a DAPI-stained nucleus and we matched the cell across photomicrographs taken using each colour independently. We also demarcated the boundaries of nuclei using the precise same fluorescent-stained and cresyl counter-stained sections. Unbiased stereology will be needed to better define the quantitative evidence reported here. Nonetheless, the evidence of bifurcating neurons was consistent across rats and, for the Re, consistent with prior studies when the mPFC was paired with the vHF. Like any neurotracer study, we cannot exclude uptake from fibres of passage, but this phenomenon has been reported to be minimal when using the CTB neurotracer protocol (Conte et al., 2009b). It seems unlikely that this confound was an issue because the highest percent of bifurcating neurons would then be expected when neurotracers were placed in closely located regions, such as the RSC when paired with the dSub; this was not the case. Conversely, locating bifurcating neurons after pairing the mPFC and caudal brain regions such as the dSub or cRSC would be the least likely to result from uptake from fibres of passage; instead, these pairings produced the highest proportion of bifurcating neurons. Additional pairings across and within the many different terminal fields of ATN efferents require evaluation in the future. It would also be important to know whether the same or different terminal neurons send reciprocal ATN afferents. The areal and laminar distribution of the axonal collaterals, and the electrophysiological properties of the bifurcating neurons relative to the non-bifurcating neurons, may provide mechanistic insights to the functional properties of this important thalamic region.

Understanding ATN function requires knowledge about how its neurons are organised and connect with other structures. The discovery of numerous bifurcating neurons in the AM suggests the need for a conceptual realignment of this region, away from a focus on segregated neural circuits and more explicitly towards concepts of control and integration of memory information across the extended memory system (Wolff and Vann, 2019). Future work will reveal the specific functional roles of the bifurcating ATN (and Re) neurons in support of memory. We suggest that bifurcating AM neurons in particular enable the explicit binding of two or more elements associated with a memory episode that convey different representations across different cortical memory structures. To learn if we are correct will demand non-conventional memory tests that go beyond the classic evidence of the ATN’s role in various aspects of spatial memory and navigation (Aggleton and Nelson, 2015; Dalrymple-Alford et al., 2015; Dupire et al., 2013; Hamilton and Dalrymple-Alford, 2021; Mathiasen et al., 2020; Nelson, 2021; Wolff et al., 2006; Wright et al., 2015).

## Materials and Methods

### Animals and housing conditions

We used 22 male Piebald Virol Glaxo cArc hooded rats, bred in the Animal Facility at the University of Canterbury and maintained in standard housing of three or four rats per opaque plastic cage (50 cm long x 30 cm wide x 23 cm high). The rats were 6-7 months at the time of surgery. Single housing was used for the 9-days of recovery following surgery, which enabled uptake of the retrograde tracers before euthanasia and histology. Food and water were available ad libitum. All procedures were approved by the University of Canterbury Animal Ethics Committee (#2018/26R).

### Surgery and infusion of retrograde neurotracers

Two Cholera Toxin Subunit B (Recombinant) Alexa Fluor™ (AF CTB) neurotracers were used. These were AF CTB 488 conjugate and AF CTB 594 conjugate (Cat# C22841 and C22842, Molecular Probes Inc., USA, supplied by Life technologies New Zealand Limited, Auckland, NZ). The tracers were used at 1% concentration (500ug in 50ul in neutral phosphate buffer), prepared by gentle rotation plus 1 sec centrifuge of the vial, in order to avoid denaturing that occurs with vortexing (Conte et al., 2009b). Vials were stored upright at 4°C for up to one week. We only used separate syringes for each tracer to prevent contamination. Each rat was injected with one tracer in one terminal region of the ATN and, in the same hemisphere, the second tracer in a second terminal region of the ATN (always the right hemisphere).

Our primary focus was to examine AM neurons projecting to the following distally-located terminal fields: (1) mPFC and dSub (n=4), (2) mPFC and vHF (n=4) and (3) mPFC and cRSC (n=3). The “mPFC and vHF” pairing has been used in previous work to identify bifurcating neurons in the Re (Hoover and Vertes, 2012; Varela et al., 2014), so this enabled a direct comparison between the AM and the rostral aspects of the Re. Given the lengthy rostro-caudal extent of the cingulate gyrus, we restricted our focus on the cRSC only, but across both areas 29b and 29c. For comparison, two additional pairings were included without the mPFC, namely (4) cRSC and dSub (n=1) and (5) cRSC and vHF (n=2). Infusions in the cRSC, however, was challenging because sometimes there was either minimal or even no sign of any CTB neurotracer. This led to the exclusion of some additional rats: mPFC and cRSC, n=2; cRSC and dSub, n=3; and cRSC and vHF, n=2. Also, one case with an intended pairing of mPFC with vHF (Case Y6) was excluded as the vHF infusion was posterior and too ventral, showing neurotracer in only the entorhinal cortex.

For infusions, rats were anaesthetized with 4% isoflurane and placed in a stereotaxic frame with atraumatic ear bars (Kopf, Tujunga, CA, USA) with the dorsal surface of the skull held in a flat position. Anaesthesia was maintained with 2% isoflurane. We infused the neurotracers via a 10 µl NanoFil syringe (blunt tip, 33-gauge needle; World Precision Instruments, Sarasota, FL, USA) at a rate of 0.05 µl / minute using a micro-infusion pump (Stoelting, Wood Dale, IL, USA). The syringe needle was lowered slowly to each coordinate and left in situ for 5 min post-infusion at each site for diffusion of the tracer. Within each target pairing, the two CTB tracers were counterbalanced across the two terminal regions. To ensure maximal coverage within each terminal region and increase uptake by ATN terminals, two x 0.6 µl (of the same tracer) infusions were made in each structure (i.e. 1.2 µl for each structure). So, an upper and a lower site was used for the mPFC; and one anterior and one posterior site was used for each of the dSub, vHF, and cRSC. For mPFC, the coordinates were: upper site (cingulate cortex, area 32D), AP + 3.0 mm from Bregma, LAT – 0.6 mm, DV – 2.2 mm; and - 3.5 mm, lower site (cingulate cortex, area 32V). For dSub: anterior, AP – 6.6 mm from bregma, LAT - 3.6 mm, DV - 3.1 mm; posterior, AP – 7.1 mm from bregma, LAT - 4.1 mm, DV - 3.3 mm. For vHF: anterior, AP - 5.28 mm from Bregma, LAT – 4.9 mm, DV - 7.4 mm; posterior, AP – 5.8 mm from Bregma, LAT - 5.2 mm, DV - 7.0 mm. And for cRSC, anterior, AP – 5.8 mm from Bregma, LAT – 0.9 mm, DV - 2.1 mm; posterior, AP - 6.6 mm from Bregma, LAT - 1.1 mm; DV - 1.8 mm.

### Histology

Rats were deeply anaesthetised with sodium pentobarbital (125mg/kg) and perfused transcardially with 200 ml of chilled saline followed by 200 ml of 4% paraformaldehyde in 0.1 M phosphate buffer solution (pH 7.4). The brains were then removed and postfixed in 4% paraformaldehyde solution overnight at 4°C. Brains were transferred into a 25% sucrose solution in phosphate buffer saline (PBS) for a minimum of 48 h before continuous 40 µm coronal sections were taken using a sliding microtome with a freezing stage (Thermofisher, Christchurch, NZ). Serial sections were collected sequentially into 24 microtubes in cryoprotectant solution (50% PBS, 25% glycerol, and 25% ethylene glycerol).

#### Fluorescence

All alternate sections (from the odd-numbered tubes) were mounted on gelatine-subbed slides and dried overnight in the dark. They were submerged in graded ethanol (10 dips in each of 70% ethanol, 95% ethanol, 100% ethanol x2), and then cleared in xylene for 5 min before mounting with antifade medium with DAPI (VectaShield® HardSet™ with DAPI, Vector Laboratories, Auckland, NZ) and a coverslip. After curing at room temperature for 15 minutes, the coverslip edges were sealed with clear nail polish. Once dry, the mounted slides were stored in light-protected boxes at 4°C. These sections were processed for fluorescence to verify the infusion site and to examine for labelling in the thalamus. The infusion site was identified in sections showing the maximum intensity of the tracer, guided by the presence of the infusion needle track.

#### Cresyl violet

After photomicrographs were taken of the fluorescent sections, at the infusion sites and in the thalamus, the same sections were re-stained for cresyl violet, thereby providing anatomical references for the fluorescent photomicrographs from the exact same sections. Slides were gently wiped with acetone to remove the nail polish and soaked in PBS overnight at room temperature to remove the coverslips. Before cresyl violet staining, the sections were processed through graded concentrations of ethanol before hydrating in distilled water (1 min) and immersion in 0.5% cresyl violet solution for 12 min. The sections were then rinsed in distilled water, dehydrated through alcohols and placed in an Acid/Alcohol differentiation bath before 100% ethanol followed by Xylene and mounting with Depex and coverslip before drying overnight.

### Photomicrographs

Sections were viewed in the dark on a Leica DM6 B upright fluorescence microscope and images captured using a DFC7000T proclass microscope camera using the LAS X imaging and analysis software (RRID:SCR_013673, Leica Microsystems). First, photomicrographs were taken using a HC PL FLUOTAR 2.5x objective (25x overall magnification) to capture the infusion sites. Regions of interest in the thalamus were identified and photomicrographs of labelled cells were taken from a series of 12 alternate sections. Photomicrographs were taken using three independent channels (red and green for the two tracers; blue for DAPI-stained nuclei) representing cells that were excited under the corresponding filter sets MCH (EX:540-580; EM:592-668), GFP (EX:450-490; EM: 500-550), and A (EX: 340-380; EM: LP 425) for Alexa Fluor® 594, Alexa Fluor® 488, and DAPI, respectively. An overlay image of the three channels combined was also taken. Photomicrographs were initially taken using the 2.5x objective to reveal the general pattern of labelled cells before using HC PL FLUOTAR 5x and then 10x objectives (50x and 100x overall magnification, respectively). When the overlaid image in the thalamus showed cells with overlap labelling of both green and red (merged into orange-yellow colour), additional photos were taken using HC PL FLUOTAR 20x and 40x (overall magnification 200x and 400x, respectively) to reveal the double-labelled cells in more detail. After the same sections were stained with cresyl violet these were photographed with the Tile Scan tool on the LAS X software.

### Cell counts

We used a semi-quantitative procedure to assess the number of single-stained and double-stained neurons using photomicrographs in regions of interest using the 20x objective. Regions of interest were subjectively identified as areas where both single-labelled and double-labelled neurons were evident. Photomicrographs were adapted for contrast adjustment within Image J (Rasband, 2011, NIH, USA). No post-imaging adjustments were made using any other software programme. We selected one representative case with well-placed infusions for each target pairing (Fig. 1, lower panels). Cell counts were taken from six representative thalamic sections, evenly spaced at about 160 µm throughout the rostro-caudal extent of the anterior thalamus (ranging from Bregma -1.20 mm to Bregma -2.16 mm). Subsequent cresyl counterstaining of the exact same sections was used to define boundaries of the thalamic nuclei for the previously acquired fluorescent photomicrographs, guided by the Paxinos and Watson atlas (7^th^ edition, 2014). Our focus was specifically the AM nucleus, which allowed us to observe bifurcating neurons in the anterior Re in the same sections (using separate photomicrographs). Single-labelled cells were counted from the individual red and green channels (corresponding to the fluorophores Alexa Fluor® 594 and Alexa Fluor® 488, respectively). However, double-labelled (i.e. bifurcating) neurons were deemed to be those showing evidence of orange-yellow colour from the overlay image of the red and green channels. Double-labelled neurons were established only when the fluorescent labelling surrounded a DAPI-stained nucleus (blue), which was checked using both 20x and 40x objectives. Also, we were careful to ensure that the bifurcating neurons could be confirmed by carefully matching the exact same cell across the independent photomicrographs taken using single red and green channels at the identical location in the section. The percentage of double-labelled cells relative to single-labelled cells for each ATN-terminal field pairing was calculated within each photomicrograph. Note, the photomicrographs did not cover the whole area of the ATN within the section under examination, and rely only on raw counts. That is, this procedure examined only the boxed region of the photomicrograph (see Figures) where photos were taken and these were used to provide an estimation across the anterior thalamic region. Cell counts were taken from photomicrographs across sections to capture a similar location within that structure at six rostral-caudal levels across different rats. To standardise counts, any cell that straddled the ventral or right edge of the photomicrograph was counted, whereas those across the dorsal or left edge were not counted. Counts were made by a researcher blind to the experimental conditions. Counts were made on a subset of the photomicrographs by a second researcher, without knowledge of the original counts, which confirmed good inter-rater reliability (R2 = 0.99, p < 0.01).

## Declaration

None of the authors have any competing interests. They acknowledge funding support from the Canterbury Medical Research Foundation, the Neurological Foundation of New Zealand, and the School of Psychology, Speech and Hearing, University of Canterbury, Christchurch, New Zealand.

## Notes

### Competing Interest Statement

The authors have declared no competing interest.

### Summary of Updates

Revised Abstract, Introduction and Discussion

